# Dynamic evolution of the heterochromatin sensing histone demethylase IBM1

**DOI:** 10.1101/2024.01.08.574644

**Authors:** Yinwen Zhang, Hosung Jang, Ziliang Luo, Yinxin Dong, Yangyang Xu, Yamini Kantamneni, Robert J. Schmitz

## Abstract

Heterochromatin constitutes a fundamental aspect of genomes that is crucial for maintaining genome stability. In flowering plants, maintenance of heterochromatin relies on a positive feedback loop involving the histone 3 lysine nine methyltransferase (H3K9), KRYPTONITE (KYP), and the DNA methyltransferase, CHROMOMETHYLASE3 (CMT3). An H3K9 demethylase, INCREASED IN BONSAI METHYLATION 1 (IBM1), has evolved to modulate the activity of KYP-CMT3 within transcribed genes. The absence of IBM1 activity results in aberrant methylation of gene bodies, which is deleterious. This study demonstrates extensive genetic and gene expression variations in *KYP*, *CMT3*, and *IBM1* within and between flowering plant species. IBM1 activity in *Arabidopsis thaliana* is uniquely regulated by the abundance of H3K9me2 in a repetitive sequence within an intron preceding the histone demethylase domain. This mechanism enables IBM1 to monitor global levels of H3K9me2. We discovered that the methylated intron is prevalent across flowering plants, however, its underlying sequence exhibits dynamic evolution. Its absence in species lacking gene body DNA methylation suggests its primary role in sensing H3K9me2 and preventing its integration into these constitutively expressed genes. Furthermore, our investigation uncovered *Arabidopsis thaliana* accessions resembling weak *ibm1* mutants, several Brassicaceae species with reduced *IBM1* expression, and a potential *IBM1* deletion. Evolution towards reduced IBM1 activity in some flowering plants could explain the frequent natural occurrence of diminished or lost CMT3 activity, as *cmt3* mutants in *A. thaliana* mitigate the deleterious effects of IBM1.

## Introduction

DNA methylation and histone H3 lysine 9 (H3K9) methylation are essential repressive chromatin modifications required for the formation of heterochromatin and the silencing of transposable elements (TEs), thereby playing a key role in maintaining genomic stability (Bannister et al., 2001; Elgin, 1996; Lachner et al., 2001; Lindroth et al., 2004; Nakayama et al., 2001). In plants, DNA methylation is observed in three different contexts: CG, CHG, and CHH (where H - A, C or T), each maintained by specific DNA methyltransferases. METHYLTRANSFERASE1 (MET1) is responsible for sustaining CG methylation through DNA replication (Finnegan et al., 1996), CHROMOMETHYLASE3 (CMT3) facilitates CHG methylation working in concert with the H3K9 methyltransferase KRYPTONITE (KYP) (Bartee et al., 2001; Du et al., 2012; Finnegan et al., 1996; Lindroth et al., 2001; Stroud et al., 2014), whereas CHH methylation is established either through the activities of CMT2 or the RNA-directed DNA methylation (RdDM) pathway (Cao and Jacobsen, 2002; Law and Jacobsen, 2010). All three contexts of DNA methylation are predominantly localized in heterochromatin and TE/repeat regions where CHG methylation (mCHG) is particularly important for reinforcing heterochromatin DNA methylation in conjunction with H3K9me2 (Du et al., 2014; Du et al., 2015; Du et al., 2012; Jackson et al., 2002; Johnson et al., 2007; Law and Jacobsen, 2010; Lindroth et al., 2001; Zhang et al., 2018). This synergy is largely due to the unique characteristics of the enzymes CMT3 and KYP, as CMT3 preferentially binds to H3K9me2, and uses it as a guide to deposit mCHG and KYP, which recognizes pre-existing DNA methylation, adds H3K9me2 (Du et al., 2014; Du et al., 2015; Du et al., 2012; Johnson et al., 2007). This interplay between CMT3 and KYP establishes a positive feedback mechanism, reinforcing the accumulation of both mCHG and H3K9me2 within heterochromatin regions (Du et al., 2015; Zhang et al., 2018).

The binding activity of KYP is not limited to mCHG, as KYP also engages with mCG prominently present in a specific group of genes classified as gene body methylated (gbM) (Bewick and Schmitz, 2017; Du et al., 2014; Johnson et al., 2007; Niederhuth et al., 2016; Tran et al., 2005). These gbM genes typically include ‘housekeeping’ genes with moderate expression, characterized by extended gene lengths, lower substitution rates (dN/dS), a higher prevalence of CWG (W = A or T, cytosines preferred by CMT3), and fewer CG dinucleotides (Bewick et al., 2019; Muyle et al., 2022; Seymour and Gaut, 2020; Takuno and Gaut, 2013; Takuno et al., 2017; Zhang et al., 2020). One prevailing hypothesis is that gbM is primarily established by CMT3, supported by the fact that the natural absence of *CMT3* in some angiosperm species is associated with the loss of gbM (Bewick et al., 2016; Bewick et al., 2017; Kiefer et al., 2019). The presence of mCG within gbM genes likely facilitates KYP binding, recruiting the CMT3-KYP heterochromatin complex and exposing these genes to silencing machinery. However, the CMT3-KYP heterochromatin feedback loop in genic regions is disrupted by the histone lysine demethylase, INCREASED IN BONSAI METHYLATION1 (IBM1), which selectively demethylates H3K9me2 in genes, thus safeguarding them from silencing (Inagaki et al., 2010; Ito et al., 2015; Miura et al., 2009; Saze et al., 2008). This protective role of IBM1 is underscored in *ibm1* mutants, which exhibit diverse phenotypic abnormalities and an accumulation of H3K9me2 and mCHG in approximately one-fifth of coding genes (Cheng et al., 2022; He et al., 2022; Inagaki et al., 2010; Ito et al., 2015; Saze et al., 2008). These affected genes in *ibm1* predominantly belong to the category of gbM genes (Zhang et al., 2021), indicating a targeted recruitment of the CMT3/KYP complex to these specific loci. The dynamic interplay between IBM1 and CMT3/KYP is important for maintaining the equilibrium between euchromatin and heterochromatin, suggesting a co-evolutionary relationship (Zhang et al., 2021). In *Arabidopsis thaliana*, the seed fertility defect and meiotic abnormalities observed in *ibm1* is rescued by knocking out *CMT3*, indicating a functional interdependence (He et al., 2022; Saze et al., 2008). Furthermore, the exclusive presence of both *IBM1* and *CMT3* in flowering plants supports the evolutionary connection between these two genes (Bewick et al., 2017).

A unique aspect of *IBM1* is its dependency on DNA and H3K9 methylation within its large 7th intron for transcriptional and post-transcriptional regulation (Rigal et al., 2012; Saze et al., 2013; Wang et al., 2013). *IBM1* is a ubiquitously expressed gene and is known to produce two distinct mRNA isoforms in *A. thaliana*. *IBM1-L*, the longer isoform, encodes a functional protein with a catalytic JmjC domain, whereas its shorter counterpart, *IBM1-S*, is non-functional without the catalytic JmjC domain (Rigal et al., 2012). Notably, the expression of these isoforms is influenced by DNA and H3K9 methylation within the *IBM1* intron. In the case of *A. thaliana* Col-0, the 7th intron of *IBM1* contains both H3K9me2 and DNA methylation, crucial for the expression of the functional *IBM1-L* isoform (Rigal et al., 2012; Saze et al., 2013; Wang et al., 2013). Full-length IBM1 has the capability to remove H3K9me2 in genic regions, including itself, suggesting that intron methylation in *IBM1* serves as a regulatory sensor of H3K9me2 by modulating the balance of its transcript isoforms (Inagaki et al., 2010; Ito et al., 2015; Rigal et al., 2012). Furthermore, our previous study revealed that certain natural *A. thaliana* accessions exhibiting increased mCHG in genic regions also show decreased intron methylation in the *IBM1* gene, along with an alteration in the *IBM1-S/IBM1-L* ratio compared to Col-0 (Zhang et al., 2020). This raises questions about the extent to which this intron methylation sensor mechanism is conserved among different *A. thaliana* accessions and across other flowering plant species and how this shapes the epigenome.

This study explores the association between intron DNA methylation of *IBM1* and its role in its own expression by surveying within and between species variation. *A. thaliana* accessions were identified that were reminiscent of weak *A. thaliana ibm1* mutants, as they possessed ectopic mCHG in a subset of genes. Furthermore, a comparative analysis of *IBM1* orthologs across 34 angiosperm species demonstrated the presence of intronic DNA methylation within its 7th intron, indicating the evolutionary conservation of the H3K9me2 sensor in flowering plants. However, the sequence underlying the methylated intron was high variably between species suggesting this heterochromatin sensing activity has a high evolutionary turnover. Moreover, our investigation into multiple Brassicaceae species suggests the coevolution of *IBM1* and *CMT3* within this family and likely all flowering plants. This is particularly evident in Brassicaceae species that lack gbM, such as *Eutrema salsugineum* and *Thlaspi arvense*, as we observed a correlation between low or absent *CMT3* expression and reduced *IBM1* expression. This association was further supported by DNA methylome data from other Brassicaceae species that have reduced/absent *IBM1* and/or *CMT3* function as well as gene body DNA methylation. Collectively, our study shows that *IBM1*, its intronic heterochromatin sensor and *CMT3* are dynamically evolving and that this shapes the genic methylation landscape in plants.

## Results

### Reduced intronic methylation of *IBM1* is associated with ectopic genic hypermethylation in natural *A. thaliana* accessions

IBM1 is a histone demethylase that removes H3K9me2 from transcribed regions in *A. thaliana* (Inagaki et al., 2010; Miura et al., 2009; Saze et al., 2008). The functional loss of *IBM1* results in an accumulation of H3K9me2 and non-CG methylation (mCHG) in a subset of gene bodies (Rigal et al., 2012; Saze et al., 2008; Ito et al., 2015; Zhang et al., 2021). The 7th intron of *IBM1* is able to sense genome-wide levels of H3K9me2, which affects *IBM1* transcription and H3K9 demethylase activity to maintain H3K9me2 levels genome wide (Rigal et al., 2012). Mutants deficient in DNA methylation, such as *met1* and *cmt3,* exhibit a decrease in the transcription of full-length *IBM1* transcripts along with a loss or reduction of DNA methylation in the *IBM1* intron region (Rigal et al., 2012). In *met1*, decrease of *IBM1* expression induces the ectopic gain of mCHG and H3K9me2 in multiple genes (Rigal et al., 2012), similar to how it occurs in *ibm1* (Inagaki et al., 2010; Ito et al., 2015; Miura et al., 2009; Saze et al., 2008; Zhang et al., 2021). Intriguingly, our previous research has identified three natural *A. thaliana* accessions exhibiting a weak *ibm1*-like phenotype, characterized by ectopic mCHG-gain in a subset of genes (Zhang et al., 2021). These accessions also showed a decrease in the DNA methylation level within *IBM1’s* 7th intron, suggesting that the heterochromatin sensor activity of *IBM1* is potentially evolving (Zhang et al., 2021).

To further explore the relationship between intron methylation of *IBM1* and ectopic mCHG in gene bodies, we used RNA-seq isoform quantification using transcriptomes from the 1,001 genomes project (n=635 with both RNA-seq and DNA methylation data, Dubin et al., 2015; Kawakatsu et al., 2016; Schmitz et al., 2013; The 1001 Genomes Consortium, 2016). Our analysis revealed that eleven *A. thaliana* accessions possess ectopic mCHG in at least 120 genes (Fig. 1A, labeled top six by name), thereby categorizing them as *ibm1*-like accessions. Notably, these *ibm1*-like accessions tend to show lower intron methylation in both CG and CHG contexts compared to the rest of the population (Fig. 1B-1C). The observed decrease in intron methylation is not due to genetic variation among the accessions, as the DNA methylation mapping coverage of the heterochromatin sensing intron was comparable across the accessions (Fig. S1A-D and Table S1). Furthermore, in accessions exhibiting low mCHG levels within the *IBM1* intron, a positive correlation (p<0.035) was observed between intron DNA methylation levels and *IBM1* gene expression (Fig. 1D). Conversely, there was a negative correlation with the proportion of short isoform *IBM1* transcripts (Fig. 1D-E and Fig. S1E-G). Consistent with this observation, the *ibm1*-like accessions tend to exhibit reduced expression of *IBM1* and an increased proportion of the *IBM1-S/IBM1-L* transcript isoforms compared to other accessions (Fig. 1F-1G). This diminished IBM1 activity likely contributed to the onset of ectopic genic methylation in these accessions. The reduced expression of functional IBM1 in these accessions raises a question regarding the plant’s potential regulatory response, specifically whether there is a compensatory downregulation of *CMT3* expression to mitigate the effects of imbalanced genic methylation. However, our investigations reveal that *CMT3* expression levels did not show significant alterations in the accessions exhibiting an *ibm1*-like molecular phenotype (Fig. S1H). Collectively, our findings indicate a robust association between the DNA methylation status of the *IBM1* intron, the ratio of the short and long transcript isoforms and the gene expression levels of *IBM1*. This relationship appears to be a significant factor influencing the widespread acquisition of mCHG in the gene bodies in a subset of *A. thaliana* accessions.

**Figure 1.**
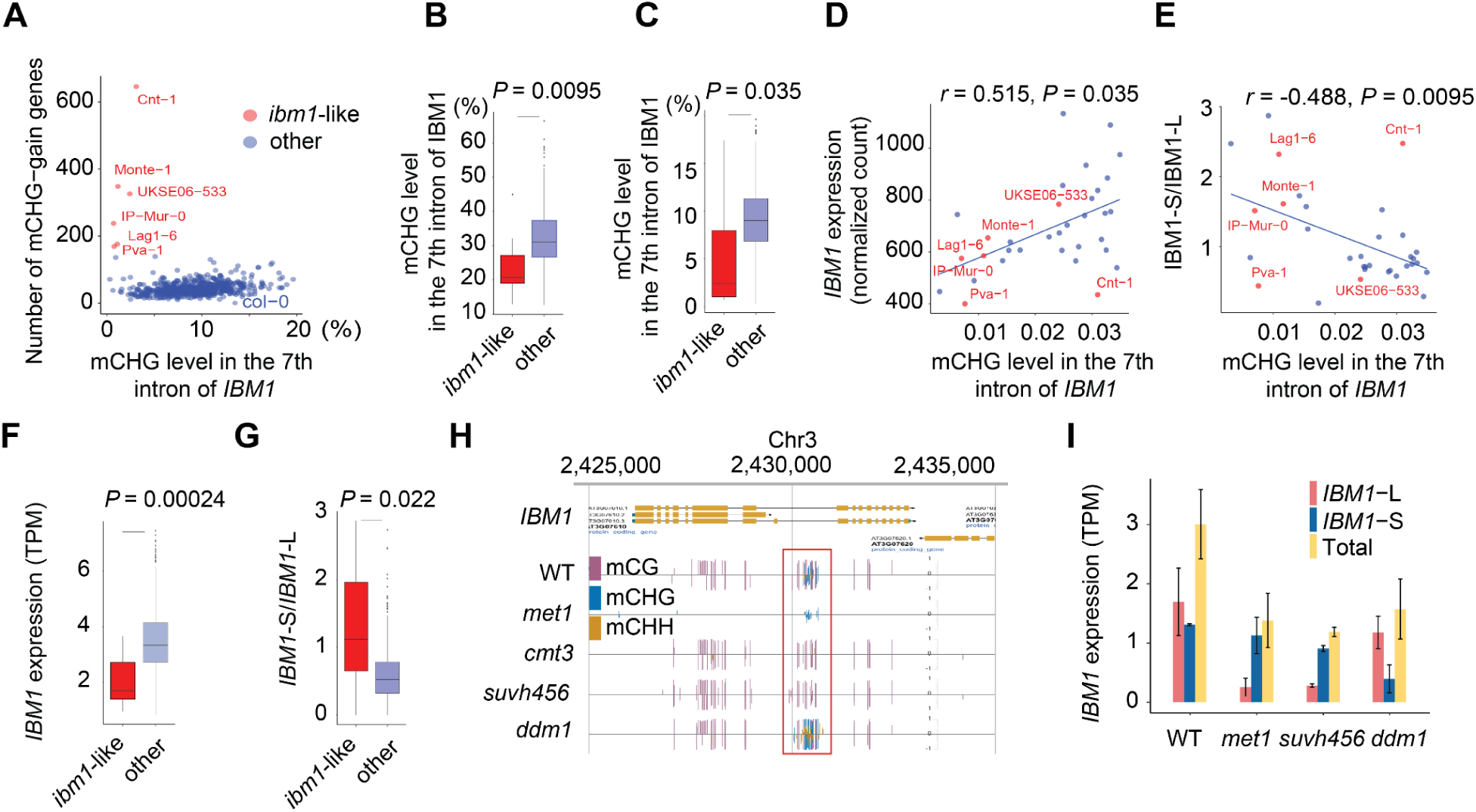
Natural variation in intron DNA methylation of *IBM1* within *A. thaliana.* **A.** The scatter plot shows the mCHG levels in IBM1’s long 7th intron against the number of genes with ectopic non-CG methylation in each accession. Accessions with a significant number (n≥130) of ectopic methylated genes are labeled by name and marked with red dots, whereas other accessions are represented with blue dots. The box plot displays the distribution of **(B)** mCG and **(C)** mCHG levels in the long intron of IBM1, comparing the top 12 accessions with the highest number of ectopic methylated genes against all other accessions. **D.** The scatter plot shows the relationship between mCHG levels in the long intron of *IBM1* and *IBM1* gene expression for accessions with low *IBM1* long intron mCHG levels (<= 0.035). The dots are colored following the same scheme as in (A)**. E.** Similar to D, this scatter plot presents the mCHG level in the long intron of *IBM1* plotted against the ratio of short to long isoform of *IBM1*. **F.** The box plot shows the distribution of *IBM1* gene expression by comparing accessions that rank in the top 12 for the highest number of ectopic methylated genes with all remaining accessions. **G.** Similar to F, the box plot shows the distribution of the ratio of short to long isoform of *IBM1* between the two groups. **H.** The browser shows the intron methylation of *IBM1* in Col-0 WT, *met1*, *cmt3*, *suvh456* and *ddm1* mutants. **I.** The bar plot shows the gene expression levels of the long and short isoforms of *IBM1*, as well as their combined expression, in *ddm1*, *met1*, *suvh456* mutants, and Col-0 WT. The isoform expressions were quantified using RNA-seq data sourced from the Saze lab (Le et al., 2020).

To validate the accuracy of RNA-seq based isoform quantification, we conducted a detailed analysis of *IBM1* isoform expression across various DNA methylation mutant lines using different data sources, including *met1, suvh456, cmt3, ibm2, ddm1,* and *drd1* (Table S2; Inagaki et al., 2017; Le et al., 2020; Ning et al., 2020; Saze et al., 2013; Zemach et al., 2013). Mutants showing a reduction in *IBM1* intron methylation, such as *suvh456, cmt3*, and *met1*, demonstrated a decrease in full-length *IBM1-L* expression level (Fig. 1H-I, Fig. S2A-C, and Table S2). In *ibm2,* where the gene encoding the enzyme crucial for *IBM1’s* proper transcription is affected (Saze et al., 2013; Wang et al. 2013), also exhibited significant reduction of the *IBM1-L* transcript (Fig. S2D). Conversely, in *ddm1* and *drd1* where the intronic methylation of *IBM1* is unaltered, the transcription of the full-length transcripts remain unaffected (Fig. 1I and Fig. S2C), indicating a direct correlation between intron DNA methylation status and the expression of functional *IBM1-L* isoform (Rigal et al., 2012).

To assess the prevalence of the role of *IBM1* intron methylation in transcription, we analyzed all *A. thaliana* genes with long introns (> 1kb). A total of 109 out of the 705 genes identified exhibit non-CG methylation within their introns (Table S3). We observed that genes with mCHG in the intron generally show a marked decrease in expression compared to those without mCHG (Fig. S2E). However, for most genes with mCHG in their introns, a reduction in methylation— ascertained by comparing *suvh456* mutants with wild-type plants—does not influence the production of full-length transcripts (Table S4). The notable exceptions to this are *IBM1* and *PPD7 (AT3G05410)*, a component of the thylakoid lumen proteome essential for the photosystem II oxygen-evolving complex in chloroplasts (Fig. S2F-G). This association was previously established in *ibm2* mutants (Duan et al., 2017). Yet, in natural accessions, the pronounced effect of intron methylation reduction on *PPD7* transcription observed in mutants does not persist. For instance, intron methylation is completely absent in the Cnt-1 accession, which surprisingly does not impact the generation of full-length transcripts. Intriguingly, minimal read mapping occurs within the intron region in Cnt-1, a long region (5.7 kb) with a non-LTR retrotransposon in the Col-0 accession (Fig. S2H-I). This suggests a possible deletion of the entire region in Cnt-1. While some accessions exhibit low read coverage, the majority demonstrate adequate coverage; yet, even with low mCHG levels, no substantial effect on the transcription process is evident in most accessions (Fig. S2I-J). This indicates that factors beyond total intron deletion, such as sequence mutations within these large introns, also may contribute to variations in transcriptional responses.

### The ectopic mCHG in genes in Cnt-1 is reduced by *IBM1* overexpression

The *A. thaliana* Cnt-1 accession stands out as an exceptional case, exhibiting the largest number of ectopic mCHG-gain genes among the natural accessions along with lower levels of intron methylation (Fig. 1A). As this phenotype resembles that of a weak *ibm1* mutant, we explored isoform quantification analysis of Cnt-1 to estimate *IBM1’s* expression level. Notably, Cnt-1 displayed a substantially lower expression level of the functional *IBM1-L* isoform in comparison to the Col-0 reference accession, with only a marginal reduction in the expression of the *IBM1-S* isoform (Fig. 2A). This result suggests that decreased expression of *IBM1-L* in Cnt-1 leads to an elevation in mCHG within these genes. Moreover, TE methylation in Cnt-1 is relatively high among all accessions (as shown in Fig. S1I and Table S1), indicating robust CMT3-KYP methylation activity. This combination of subdued IBM1 activity and vigorous CMT-KYP methylation may account for the high number of mCHG-gain genes in Cnt-1 and supports that this heterochromatin sensing mechanism is dynamic in populations to shape the epigenome.

**Figure. 2.**
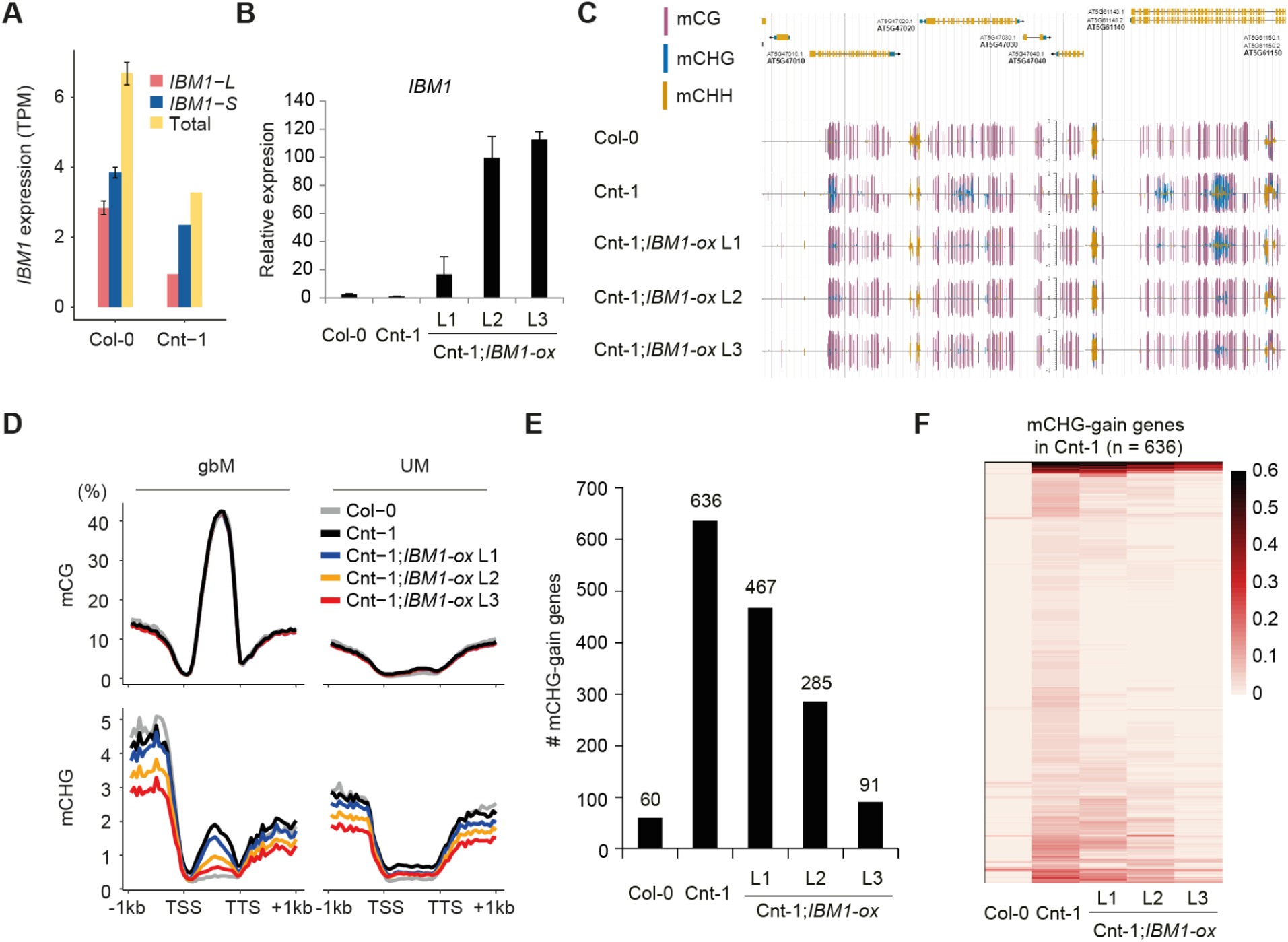
The ectopic mCHG in genes in Cnt-1 is reduced by *IBM1* overexpression. **A.** Expression level of *IBM1-L* and *IBM1-S* in Col-0 and Cnt-1. **B.** Expression level of *IBM1* in each transgenic plant. Primers were designed to target the conserved region of *IBM1* between Col-0 and Cnt-1. **C.** A genome browser view shows that ectopic mCHG in genes in Cnt-1 is reduced by introducing the *IBM1* transgene. **D.** Metaplots show the average changes to mCHG over gbM and UM genes in *IBM1* transgenic lines of Cnt-1. **E.** The number of mCHG-gain genes is reduced in *IBM1* transgenic lines compared to Cnt-1. **F.** Heatmaps show reduction of mCHG on genes in *IBM1* transgenic lines of Cnt-1. The 636 mCHG-gain genes from Fig. 2E are shown.

Next, we examined the effect of restoring *IBM1* expression in the Cnt-1 accession. The *UBQ10* cis-regulatory sequences were used to express the Col-0 *IBM1* coding sequence in Cnt-1, resulting in the generation of three independent transgenic lines. These lines exhibited a significant increase in *IBM1* expression, ranging from 20-100-fold higher than the control (Fig. 2B). All three of the *IBM1*-ox lines displayed a decrease in mCHG in a subset of gene bodies compared to the Cnt-1 control (Fig. 2C and D), indicating that overexpression of *IBM1* reduces ectopic mCHG in gene bodies presumably by decreasing H3K9me2, although this was not tested. This reduction was further supported by a decrease in the number of mCHG-gain genes when compared to the Cnt-1 control (Fig. 2E). A heatmap analysis of 636 mCHG-gain genes from Cnt-1 revealed a substantial decrease in mCHG levels in the *IBM1*-ox lines (Fig. 2F). Considering the conservation of *IBM1* coding sequences between Col-0 and Cnt-1 (99.61% sequence identity), these findings collectively indicate that rescuing *IBM1* expression in Cnt-1 significantly reduces the aberrant mCHG accumulation in the gbM genes.

### Methylation of the intronic heterochromatin sensor is a common feature in *IBM1* orthologs across flowering plants

We noted a heightened mutation frequency within the methylated intron of *IBM1* compared to the unmethylated non-coding regions and the coding sequences of *IBM1* across *A. thaliana* populations (Fig. S3A) as well as at the species level (Fig. S3B) (Table S5). Given these observations, further exploration into the variability of *IBM1* intronic DNA methylation and the conservation of its regulatory role across different species presents an intriguing line of inquiry. To investigate the patterns of intronic DNA methylation of *IBM1* across angiosperm species, we first identified the orthologous genes of *IBM1* in various plant species. This was achieved through the construction of a Maximum Likelihood (ML) gene tree, using 1,516 JmjC family homologous genes derived from 597 different species (see Methods and Table S6). The orthologs of *IBM1* were defined as those genes that clustered within the same clade as *A. thaliana’s IBM1*, specifically clade 5, as shown in Fig. S3C-E.

We analyzed the DNA methylation patterns on *IBM1* orthologous genes in 34 plant species for which DNA methylome data were available (Table S7; Niederhuth et al., 2016). This set includes *A. trichopoda,* the most basal species in our study, which was used as an outgroup. Our investigation focused on identifying introns within these *IBM1* orthologs that exhibit significantly higher levels of mCHG methylation compared to the species-specific background genic methylation levels (Table S8). Our findings reveal that intronic CHG methylation is prevalent in *IBM1* orthologous genes across both eudicots and monocots (Fig. 3A-B, Fig. S4). Among the 65 *IBM1* orthologs analyzed, which include species with multiple gene copies, 27 *IBM1* orthologs were significantly enriched for mCHG within their introns (Figure 3C).

**Figure 3.**
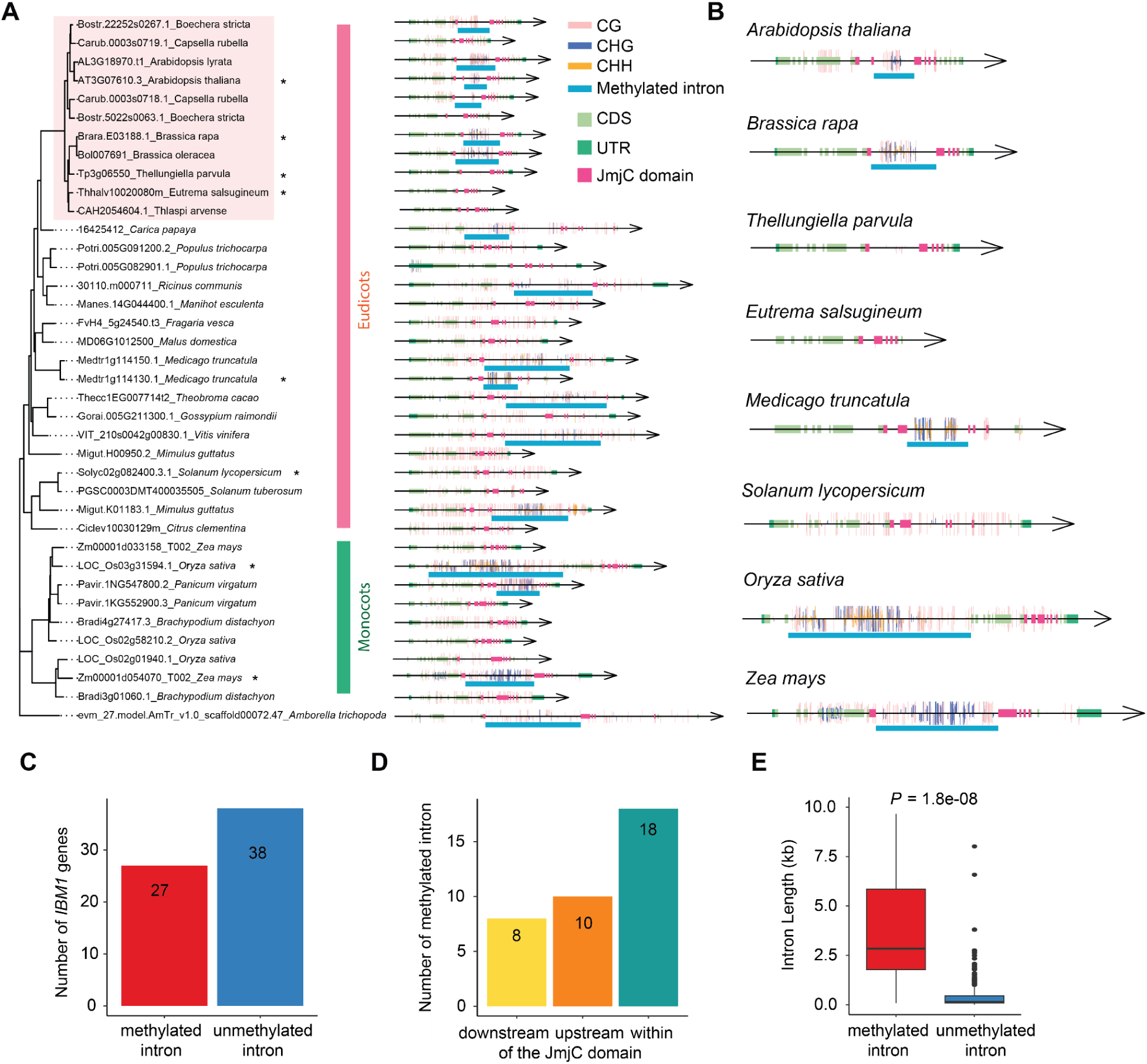
Intron methylation is widely present in *IBM1* homologous genes of angiosperms. **A.** The maximum likelihood gene tree displayed here represents the *IBM1* orthologs from 34 angiosperm plants that have DNA methylome data. Due to space constraints, this is a truncated version of the tree; the complete version is available in Fig. S4. The left panel presents the gene tree, while the right panel depicts the corresponding gene structures. UTRs, CDS, and the JmjC domain are highlighted in different colors, as indicated in the legend, with introns represented by thin black lines. The locations of methylated sites on the gene sequences are marked with vertical bars, with distinct colors denoting the different methylation contexts: CG, CHG, and CHH. **B.** Zoomed-in version of A. A few species were enlarged to highlight the intron methylation. **C.** The bar plot shows the number of genes out of the 65 *IBM1* orthologs investigated that are either with or without mCHG in their introns. To assess the significance of mCHG levels within these introns, a binomial test was applied. **D.** The bar plot displays the count of introns with mCHG relative to their position within, before, or after the JmjC domain. It should be noted that a single *IBM1* ortholog may contain multiple methylated introns. **E.** The boxplot shows the distribution of intron length between introns that are covered by mCHG and introns that are not covered by mCHG.

Within the Brassicaceae family, intronic DNA methylation is present in the *IBM1* orthologs in six out of nine species, encompassing all species from the *A. thaliana* subclade. The intronic methylation observed in these *IBM1* orthologs is predominantly found within the JmjC domain, a pattern that is consistent with what has been observed in *A. thaliana*. A varied pattern emerges in the *E. salsugineum* subclade: *B. rapa* and *B. oleracea* exhibit significant mCHG enrichment in the long intron region within the JmjC domain, whereas *T. parvula* and the species with shorter introns, *E. salsugineum* and *T. arvense*, show a loss of mCHG. This is noteworthy, as the species with no or reduced DNA methylation in the *IBM1* intron region containing the heterochromatin sensor are the species that have lost gbM (Bewick et al., 2016; Galanti et al., 2022). For species outside the Brassicaceae family, the patterns of intronic methylation in *IBM1* orthologs are more variable and not strictly confined to the JmjC domain. DNA methylation is observed in regions both upstream and downstream of the JmjC domain (Fig. 3A, 3D, Fig. S4), and this variability in DNA methylation location does not appear to be conserved in species from the same family. For instance, in the Fabaceae family, the *IBM1* orthologs in *L. japonicus* and *G. max* exhibit intronic methylation after the JmjC domain, whereas *M. truncatula* from the same family shows DNA methylation within the JmjC domain. A similar pattern is noted in the Poaceae family: an *IBM1* ortholog in *P. virgatum* has intronic DNA methylation within the JmjC domain, whereas in *O. sativa*, it is located before the JmjC domain. The mCHG introns are typically more enriched in longer introns (Fig. 3E). For instance, a gene copy in *M. esculenta*, with three successive large introns (7.3kb, 3.6kb, 9.6kb), and another in *T. cacao*, with a 27.2 kb intron, both exhibit significant DNA methylation enrichment within these intron regions (Fig. S4). These data support that many of these species likely use a similar mechanism to *A. thaliana* to sense heterochromatin content given the persistence of intron methylation within the JmjC domain of *IBM1*.

### The DNA sequence underlying the heterochromatin sensor of *IBM1* is dynamically evolving

In *A. thaliana,* DNA methylation in the 7th intron is present within a 150 bp repeat sequence that shows high similarity to *YCF1*, a gene encoded by the chloroplast genome. Organellar sequences are often silenced by DNA methylation upon integration into the nuclear genome. In the *IBM1* ortholog from *A. lyrata*, the DNA methylated region in the 7th intron is larger than that in *A. thaliana*, as it arises from two insertions of simple repeat elements approximately 200 bp upstream and 50 bp downstream of the sequence aligned with the DNA methylated intron of *A. thaliana* (Fig. 4A). Intriguingly, this *YCF1*-like fragment is identified in *A. lyrata,* but not other Brassicaceae species, suggesting significant evolutionary divergence.

**Figure 4.**
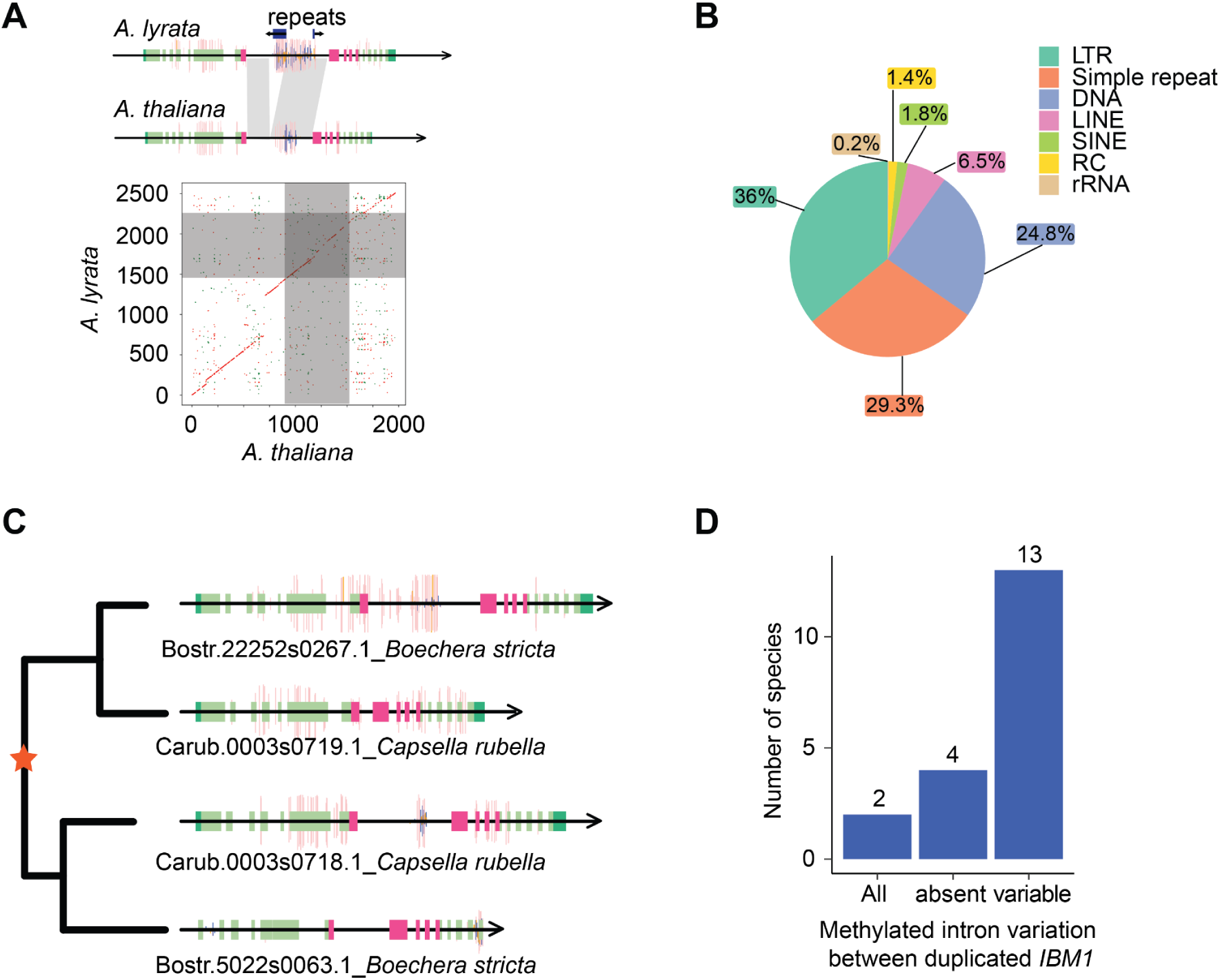
The sequence underlying the methylated intron of *IBM1* is highly divergent between species. **A.** The plot illustrates the sequence comparison between *A. lyrata* and *A. thaliana,* which reveals an insertion of approximately 500 bp in the intron of *A. lyrata*. This inserted fragment comprises repeat sequences that are methylated. **B.** The pie chart shows the various types of DNA repeats found in the introns covered by mCHG across the 65 *IBM1* orthologs. **C.** The plot shows a distinct example of differing intronic methylation patterns between duplicated *IBM1* genes in *B. stricta* and *C. rubella.* **D.** The bar plot displays the distribution of intronic methylation variation among 19 species that have duplicated *IBM1* gene copies. The categories are defined as follows: ’All’ indicates that every *IBM1* gene copy in a species has a methylated intron, ’No’ signifies that no copies of the *IBM1* genes in a species have methylated introns, and ’Varied’ denotes species where the methylation status differs among the *IBM1* gene copies.

It is likely the heterochromatin sensing ability of *IBM1* is conserved throughout many flowering plants given the presence of mCHG in intron(s) within the JmjC domain of many species (Fig. 3A). However, the sequence underlying the methylated introns is not conserved within species (Fig. 4B). We found numerous unique simple repeats and transposon repeats showing that sequence evolution within the intronic heterochromatin sensing region of *IBM1* is quite dynamic between species (Fig. 4B). For species that have a candidate heterochromatin sensor in an *IBM1* intron, the actual sequence that is methylated isn’t as important as the methylation event itself. The fact that the sequence is continually changing between species, but remains methylated in many flowering plants suggests that there could be constant gains and loss of *IBM’s* ability to sense H3K9me2 levels over evolutionary time.

In fact, DNA methylation patterns in the intronic region of *IBM1* orthologs not only varied among different species, but also among duplicated gene copies within the same species. For instance, a duplication event happened just before the divergence of *B. stricta* and *C. rubella*, resulting in each species possessing two *IBM1* gene copies, and DNA methylation profiles also varied between the two gene copies (Fig. 4C). Approximately 20% of species in this clade possess duplicated *IBM1* genes (Fig. S3F) and out of the 34 species for which there is DNA methylation data, 19 species contain duplicated *IBM1* genes, and over half exhibit DNA methylation pattern divergence between the duplicates (Fig. 4D). This indicates that upon duplication the heterochromatin sensor activity is not retained in duplicate copies.

### Gene body DNA methylation is frequently lost in Brassicaceae species due to reduction of CMT3 or IBM1 activity

Previous studies have shown that loss of *CMT3* or reduced CMT3 activity is associated with a loss or a reduction of gbM (Bewick et al., 2016; Kiefer et al., 2019). Importantly, the loss of *CMT3* has occurred multiple independent times suggesting that this is an evolving process (Bewick et al., 2016; Kiefer et al., 2019). One possible reason for the loss of *CMT3* or a reduction in its activity could be due to reduced function of IBM1. Loss of IBM1 activity in *A. thaliana* leads to reduced expression of gbM genes and eventually lethality (Saze et al., 2008. However, this is rescued by reduced activity or a complete loss of CMT3 (He et al., 2022; Saze et al., 2008). To explore this possibility further, we analyzed transcriptomes and DNA methylome data from various Brassicaceae species, specifically focusing on *IBM1, CMT3* and *KYP* expression and the prevalence of gbM (Fig. 5A). In *E. salsugineum*, a noticeable decrease in *IBM1* expression accompanies the loss of *CMT3*, coinciding with a substantial reduction in gbM. A similar pattern emerges in *T. arvense*, closely related to *E. salsugineum*. Although *T. arvense* retains *CMT3*, both *IBM1* and *CMT3* exhibit low expression levels, correlating with an absence of gbM. This trend persists within the Brassicaceae subclade; species like *B. rapa* and *B. oleracea*, showing diminished *CMT3* and *IBM1* expression, also display a marked decrease in gbM genes (Fig. 5A and Table S9). This is consistent with our previous results showing *CMT3* in some Brassicaceae is under relaxed selective constraints (Bewick et al., 2017).

**Figure 5.**
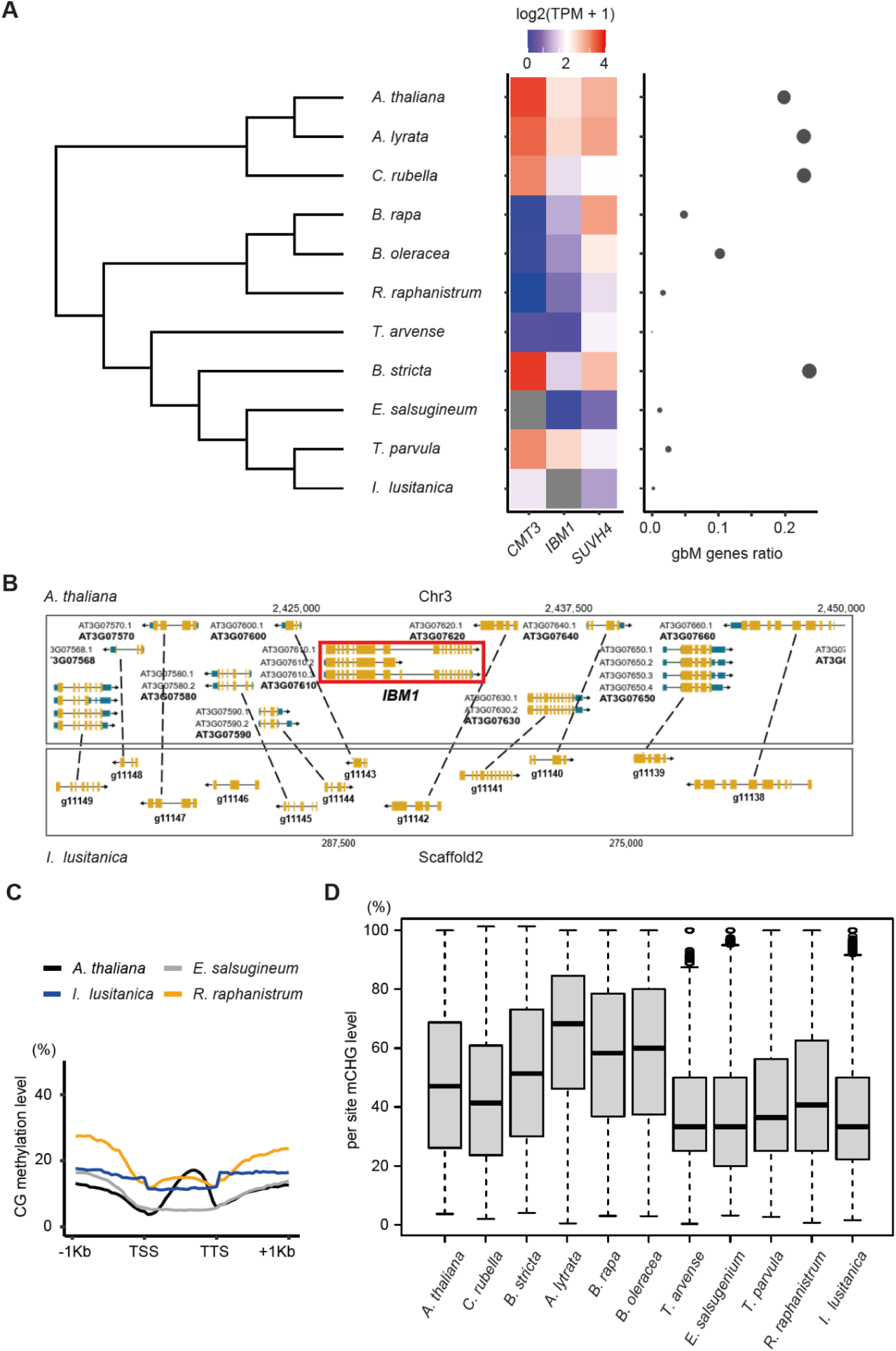
Frequent expression and genetic variation of *IBM1* and *CMT3* in Brassicaceae. **A.** The heatmap on the left panel displays the expression levels of *CMT3, SUVH4*, and *IBM1* in Brassicaceae species. On the right panel, the scatter plot shows the gbM gene ratio for each species, which is calculated as the number of gbM genes divided by the total number of genes where DNA methylation could be measured in that species. The size of the points in the scatter plot is proportional to the gbM gene ratio. B. A genome browser view of a region of synteny between *I. lusitanica* and *A. thaliana* depicting the absence of *IBM1.* **C.** mCG metaplots over all genes in newly analyzed Brassicaceae species. **D.** Box plots showing per-site mCHG level across the genome, which reflects the enzyme activity of CMT3.

By mining publicly available reference Brassicaceae genomes, we identified two additional species (*Raphanus raphanistrum* and *Isatis lusitanica*) with potential loss or truncations of *IBM1* or *CMT3* (Figure S5A-B). However, the quality of these reference genomes were quite variable, requiring additional experiments to validate the loss or truncations of *IBM1* or *CMT3.* We used transcriptome assembly as well as PCR of genomic DNA using universal primers to extract full-length *IBM1* or *CMT3* loci (Table S10). We were unable to identify *IBM1* in *I. lusitanica* suggesting it might be lost in this species. There is clear evidence that it is deleted within a region of synteny with *A. thaliana* (Fig. 5B). We successfully identified *CMT3* sequences across both species, although *I. lusitanica* exhibited a small N-terminal deletion in *CMT3* (Figure S5C, Table S11). Next, we estimated the expression level of *IBM1, CMT3* and *KYP* across these species and evaluated the presence of gbM genes (Fig. 5A). The number of potential gbM detected in *I. lusitanica* and *R. raphanistrum* was as low as *E. salsugineum* (Fig. 5A and B) indicating it has been lost in these species. In *R. raphanistrum*, *CMT3* expression was absent, coupled with a significant reduction in *IBM1* expression. In *I. lusitanica*, in which we couldn’t identify *IBM1*, there were relatively lower *CMT3* expression levels compared to Brassicaceae species with high numbers of gbM genes (Fig. 5A).

We hypothesized that even though some of these species have *CMT3* that its expression or enzymatic activity might be reduced or absent consistent with the reduction of gbM (Fig. 5C). We tested this by measuring mCHG levels and mCHG symmetry, both of which should be present if CMT3 activity is functional. Genome-wide per-site mCHG levels were measured across species, revealing *R. raphanistrum* and *I. lusitanica* with low mCHG levels, similar to gbM-absent species such as *E. salsugineum* (Fig. 5D). We also assessed the methylation symmetry at CWG sites (W = A or T) preferentially targeted by CMT3. Furthermore, *R. raphanistrum* and *I. lusitanica* exhibited a predominantly asymmetric mCWG pattern, aligning with the pattern observed in *E. salsugineum*, which lacks *CMT3* (Figure S6).

## Discussion

The discovery of heterochromatin sensing activity associated with *IBM1* in *A. thaliana* supports that mechanisms have evolved to buffer heterochromatin abundance in flowering plant genomes (Rigal et al., 2012; Zhang et al., 2020). This discovery parallels the DNA methylation sensing activity in *A. thaliana* associated with the DNA demethylase *ROS1* (Lei et al., 2015; Williams et al., 2015). Epigenome homeostasis is an emerging property associated with chromatin regulation and genome evolution (Murphy and Berger, 2023; Williams and Gehring, 2020). The existence of these epigenome sensors suggests that mechanisms are in place to cope with wholescale changes to H3K9me2 or DNA methylation patterns in plants that could occur due to epigenome shock, whole genome duplication and/or hybridization events among other possibilities.

This study explored the evolution of heterochromatin sensor activity within an intron that disrupts IBM1 activity if it is not properly spliced in *A. thaliana*. We identified both within and between species variation of IBM1 activity. Some *A. thaliana* accessions were identified that had ectopic mCHG in gene bodies, which was rescued by ectopic expression of *IBM1.* Variation in the related histone lysine demethylase *JMJ26* was recently discovered in a genome-wide association study for mCHG variation in transposons (Sasaki et al., 2022). This provides precedent for natural variation of histone demethylase activity in shaping plant epigenomes.

It’s unknown whether *IBM1’s* heterochromatin sensing ability extends outside of *A. thaliana.* DNA methylation of the intron within the JmjC domain of *IBM1* was common throughout flowering plants with notable exceptions in species that do not have gbM such as *E. salsugineum* and *T. arvense.* This could suggest that once gbM is lost there is no need for *IBM1* sensing activity, as ectopic H3K9me2/mCHG can no longer be recruited to these genes without CMT3 activity. We also identified numerous duplicate *IBM1* copies where one copy retained the methylated heterochromatin sensor intron and the other did not. This could indicate that dosage associated with sensing activity is important to maintain heterochromatin abundance in these species. We also observed that the sequence underlying the methylated heterochromatin sensor intron of *IBM1* was dynamically evolving between species even though heterochromatin sensor activity is likely retained. This suggests that there is a dynamic interplay between the evolution of heterochromatin abundance and the activity of IBM1 and CMT3. Identifying the triggers of this highly evolving process will be important for understanding how heterochromatin abundance is modified during flowering plant genome evolution.

It is curious that there are repeated occurrences of loss or reduced activity of CMT3 along with reduced or loss of gbM in some plant genomes. Why are there so many examples of deletion and/or loss of CMT3 activity? One possibility could be that there are genome-wide events that disrupt epigenome homeostasis, such that IBM1 no longer efficiently removes H3K9me2 from gbM genes. This would likely lead to a rapid loss of fitness due to decreased expression of many ‘housekeeping genes’. However, secondary site mutations in CMT3 would eliminate this silencing effect. We have identified numerous examples in this study where there is support for this model. For example, *E. salsugineum* has lost *CMT3* and has reduced expression of *IBM1*. We even identified a species that has potentially lost *IBM1* (*I. lusitanica*). Even though *I. lusitanica* has *CMT3*, mCHG levels and symmetry analysis of mCHG shows that it is not functional supporting it could be a natural double mutant of *IBM1* and *CMT3*. Higher quality genome sequence efforts will be required to confirm the loss of *IBM1*. Regardless, this species has lost CMT3 activity and gbM. Future studies are also needed to evaluate the extent to which *IBM1’s* heterochromatin sensor is functional across flowering plants. This is difficult to assay, given its discovery was dependent on the use of DNA methylation mutants in *A. thaliana,* which are not well tolerated in many flowering plants. Continual exploration of the evolution of *IBM1* and *CMT3* in newly released flowering plant genomes, especially within the Brassicaceae, will deepen our understanding of plant epigenome homeostasis.

## Acknowledgements

We thank Frank Johannes for critical feedback on this manuscript. R.J.S. acknowledges support from the National Science Foundation (MCB-2242696).

## Methods

### WGBS and RNA-seq data acquisition

We obtained whole genome bisulfite sequencing (WGBS) and RNA-seq data for natural accessions of *A. thaliana* from the 1,001 Genomes Project (Dubin et al., 2015; Kawakatsu et al., 2016; Schmitz et al., 2013; The 1001 Genomes Consortium, 2016). The WGBS and RNA-seq data for *A. thaliana* mutants were obtained from NCBI (Inagaki et al., 2017; Le et al., 2020; Ning et al., 2020; Saze et al., 2013; Zemach et al., 2013), with specific details provided in Table S2. Additionally, we acquired WGBS and RNA-seq data for 34 angiosperm species from NCBI (Amborella Genome, 2013; Bewick et al., 2016; Niederhuth et al., 2016; Stroud et al., 2013; Zhong et al., 2013; Zhou et al., 2020a; Zhou et al., 2020b), with the sources detailed in Table S7. Furthermore, we generated WGBS and RNA-seq data for 2 Brassicaceae family species, and both the sequencing and processed data are available from GEO under the accession number GEOXXX.

### Plant materials and transgenic line analysis

The *IBM1* coding sequence from Col-0 was driven by the *UBQ10* promoter. The construct was transformed into the *A. thaliana* Cnt-1 accession by *Agrobacterium*-mediated flower dipping method (Clough and Bent, 1998). Three transgenic lines were selected and RNA was extracted to estimate *IBM1* expression. The genomic DNA from those lines were subjected to WGBS analysis. For the DNA methylation and RNA-seq analyses of the four Brassicaceae species, namely *Physaria fendleri*, *Biscutella auriculata, Raphanus raphanistrum*, and *Isatis lusitanica*, the plants were grown under identical conditions to those used for *A. thaliana*. Leaf tissues were collected from each species at the 8-week-old growth stage for subsequent analyses.

### Whole genome bisulfite sequencing library preparation

Libraries were prepared following the MethylC-seq protocol (Urich et al., 2015). Briefly, genomic DNA was isolated from leaf tissues using the DNeasy Plant Mini Kit (Qiagen). Subsequently, genomic DNA was sonicated to achieve 200 bp fragments, and then end-repair was performed by the End-It DNA End-Repair Kit (Epicentre). This end-repaired DNA was subjected to A-tailing using the Klenow 3′–5′ exo− enzyme (New England Biolabs). The subsequent step involved the ligation of methylated adapters to the A-tailed DNA, using T4 DNA Ligase (New England Biolabs). Following adapter ligation, the DNA was bisulfite converted with the EZ DNA Methylation-Gold Kit. Finally, the library was amplified using KAPA HiFi Uracil + Readymix Polymerase (Roche).

### DNA methylation analysis

WGBS data were processed using Methylpy (Schultz et al., 2015), following the methodology outlined in reference (Zhang et al., 2021). Initially, read quality filtering and adapter trimming were conducted using Cutadapt v1.9.dev1. The qualified reads were then aligned to the species-specific reference genome using Bowtie 2.2.4 (Langmead and Salzberg, 2012), ensuring that only uniquely aligned and non-clonal reads were retained. The genome assembly version of 34 species used for mapping are provided in Table S7. To calculate the non-conversion rate of unmodified cytosines in the sodium bisulfite reaction, a fully unmethylated sequence, chloroplast or lambda (see unmethylated sequence used for each species in Table S7), was used as a control. A binomial test, requiring a minimum coverage of three reads, was used to determine the DNA methylation status of cytosines.

To determine genes with gene body methylation (gbM) in each of the 34 species, we counted the number of methylated and total cytosines for each methylation context (CG, CHG, and CHH) within the coding regions of primary transcripts for each gene. We then calculated the percentage of methylated sites for each context across all coding regions in each species. This percentage served as the background probability of methylation at a single site within coding sequences (CDS). Using this background probability, along with the total counts of cytosines and methylated cytosines on CDS, we calculated p-values based on a binomial distribution. These p-values represent the cumulative probability of observing a greater number of methylated cytosines in a given gene than expected by chance. Subsequently, we adjusted the p-values using the Benjamini–Hochberg False Discovery Rate (FDR) method to compute q-values. A gene was classified as having gbM if it had reads mapping to at least 20 CG sites, a q-value less than 0.05 for mCG, and q-values greater than 0.95 for both mCHG and mCHH. To calculate the gbM ratio, the total number of gbM genes was divided by the overall count of genes with adequate coverage in each species. This process of calculating gene coverage required counting CG sites within each gene, selecting those genes where at least 40% of the CG sites had a minimum coverage of three reads or more.

In natural *A. thaliana* accessions, we identified ectopic non-CG genes characterized by high non-CG methylation (mCHG and/or mCHH with a q-value less than 0.05) in certain accessions, while typically existing as gbM genes in over 90% of all accessions. Specifically in Cnt-1, which has the highest number of ectopic mCHG in genes, we compared the mCHG levels of these genes between the wild type Cnt-1 and an *IBM1* overexpression transgenic line in Cnt-1 (*IBM1-ox*). The average DNA methylation ratio of each gene was calculated, and the number of the gbM genes that gained more than 2% mCHG (> 0.02) was plotted. The DNA methylation ratio of the mCHG-gain gbM genes in Cnt-1 (n = 636) were plotted as heatmaps.

Using binomial tests, we identified introns enriched with mCHG in *IBM1* of different species. For this, we counted the methylated and total cytosines for each methylation context (CG, CHG, and CHH) specifically within the 7th intron of *IBM1*. We then calculated the percentage of methylated sites for each context in all introns for each species. This species-specific percentage served as the background probability. With this background probability and the total counts of cytosines and methylated cytosines in the *IBM1* intron region, we computed p-values using a binomial distribution. These p-values were then adjusted using the Benjamini–Hochberg False Discovery Rate (FDR) method to derive q-values. An intron was classified as mCHG-enriched if its mCHG q-value was less than 0.05.

In the genic metaplots for *A. thaliana*, we divided the gene body into 20 equal windows. Similarly, the 1,000 base pairs upstream and downstream of the gene were each divided into 20 windows. Within each window, we calculated the weighted DNA methylation (Schultz et al., 2012). Subsequently, we computed the average weighted methylation for each window across all genes. These average values were then plotted using R to create the metaplots.

### Per site methylation and symmetry analysis

The methylation ratios at individual CHG sites were calculated under the condition that each CHG site exhibited a minimum read coverage of three. Additionally, a CHG site was included in the analysis only if at least one CHG site was identified as being methylated. For the analysis of methylation symmetry, we selectively focused on CWG sites to exclude the influence of MET1’s activity on CCG sites. Both strands of CWG sites were required to have a minimum coverage of at least three reads, and at least one CWG site was identified as being methylated.

### Isoform quantification for RNA-seq data

Quality filtering and adapter trimming of the RNA-seq reads were conducted using Trimmomatic v0.33 (Bolger et al., 2014), using default parameters. For each species, the processed reads were then aligned to their respective transcriptome fasta files using Kallisto v0.50.0 (Bray et al., 2016) for transcript quantification. Specifically in *A. thaliana*, to quantify the expression of short transcripts in genes with long introns, we generated truncated versions of these transcripts, comprising only the UTR and CDS regions preceding the long intron. These truncated transcripts were subsequently incorporated into the transcriptome files used for mapping. Similarly, for other Brassicaceae species with long introns in *IBM1*, we created and added shortened versions of *IBM1* transcripts into their respective transcriptome files for precise quantification of the short isoform expression of *IBM1*.

### Identification of IBM1 and CMT3 in the Brassicaceae species

The orthologs of *IBM1* and *CMT3* across four Brassicaceae species were identified using OrthoFinder (Emms and Kelly 2019), and the extracted coding sequences were used for phylogenetic analysis using MEGA (Tamura et al., 2021). The *de novo* transcriptome assembly was performed using Trinity (Grabherr et al. 2011). The presence/absence of the *IBM1* and *CMT3* transcripts were identified by BLASTing the *A. thaliana* gene sequence against the assembled transcriptome. To amplify the *IBM1* and *CMT3* gene from the genomic DNA, the degenerate primers were designed (Table S10), and the genes were PCR-amplified using Q5® High-Fidelity 2X Master Mix (New England Biolabs).

### Homologous genes identification and phylogeny analysis

We extracted the JmjC gene family from the One Thousand Plants (1KP) Consortium’s orthogroupings (One Thousand Plant Transcriptomes, 2019), using the *A. thaliana IBM1* gene identifier (AT3G07610). The 1KP Consortium identified a single orthogroup that encompasses the IBM1 proteins, along with three other JmjC family genes (JMJ27, JMJ26, JMJ29) from *A. thaliana*, with a total of 9,258 protein sequences. The corresponding coding sequences (CDS) for these genes were also obtained from the 1KP Consortium. We expanded our dataset to include sequences from 69 species not covered in 1KP, including 45 from the Brassicaceae family. Their annotated CDS and protein sequences were obtained from Phytozome or EBI, with specific data sources detailed in Supplementary Table S9. For these species not included in iKP orthogroup, protein sequences showing reciprocal best BLAST hits with *A. thaliana* JMJ27 (AT4G00990), JMJ26 (AT1G11950), JMJ29 (AT1G62310), and IBM1 (AT3G07610) were added in the orthogroup. In total, the JmjC gene family included 9,521 sequences from 1,130 species. Then, according to Interproscan, sequences were retained if they included the same PFAM domain (JmjC domain PF02373) or ProSiteProfiles domain (JmjC domain profile PS51184) as *A. thaliana*. These filtered sequences included 1,528 sequences from 602 species, from which 10 sequences were excluded due to discrepancies between their protein and CDS sequences in terms of codon-to-amino acid conversion.

Then, to estimate the JmjC gene tree, a protein alignment was carried out using Pasta (Mirarab et al., 2015) with the default setting. The resulting alignment was back-translated using the coding sequence into an in-frame codon alignment. Then, Gblocks was used to retain only conserved codons with default settings, but allowing for a 50% gapped position. This conserved codon alignment then served as the input for phylogenetic estimation using RAxML (Stamatakis, 2014), which included 500 rapid bootstrap replicates. The generated tree was rooted at the green algae clade and subsequently edited using the R package ggtree. From this comprehensive gene tree, a subtree comprising *IBM1* orthologous genes from 34 species with available methylome data was extracted using ggtree.

### Identification of introns where decreased mCHG correlates with full-length transcripts in A. thaliana

For each gene featuring a long intron (≥1kb) enriched with mCHG, we determined the average expression of both short and long isoforms across replicates, conducting these calculations independently for wild-type and *suvh456* samples. Subsequently, we identified genes exhibiting a difference greater than 2 in the short-to-long isoform ratio between *suvh456* mutants and wild type.

### Calculation of SNP mutation rates

To determine the SNP density within the *A. thaliana* population, we divided the SNP genotype data of *IBM1* genes (sourced from the 1,001 Genomes Project) into three groups: SNPs in the CDS region, SNPs within the methylated 7th intron, and SNPs in other intron regions not covered by methylation. We then calculated the SNP density and Tajima’s D values separately for each of these SNP matrices. Tajima’s D values were calculated by R package PopGenome while SNP density was calculated as the proportion of sites with SNP in each of three groups. To calculate the nucleotide mutation rate between *A. thaliana* and *A. lyrata*, we first generated separate sequence alignments for the coding sequences (CDS) and intron regions of both species. The intron regions were further categorized into methylated regions and non-methylated regions. For each of these three groups—CDS, methylated introns, and non-methylated introns—we counted the number of identical nucleotides shared between the two species. The nucleotide difference for each group was then determined by calculating 1 minus the proportion of identical bases.

### Annotation of repeats and motifs in IBM1 introns across various species

The DNA sequences of the introns for each of the 34 species were extracted from their respective genome assemblies using GFF annotations. RepeatMasker (Tarailo-Graovac and Chen, 2009) was then used to identify various types of transposable elements, utilizing Repbase (Bao et al., 2015) as the reference library.

---

**Figure S1.**
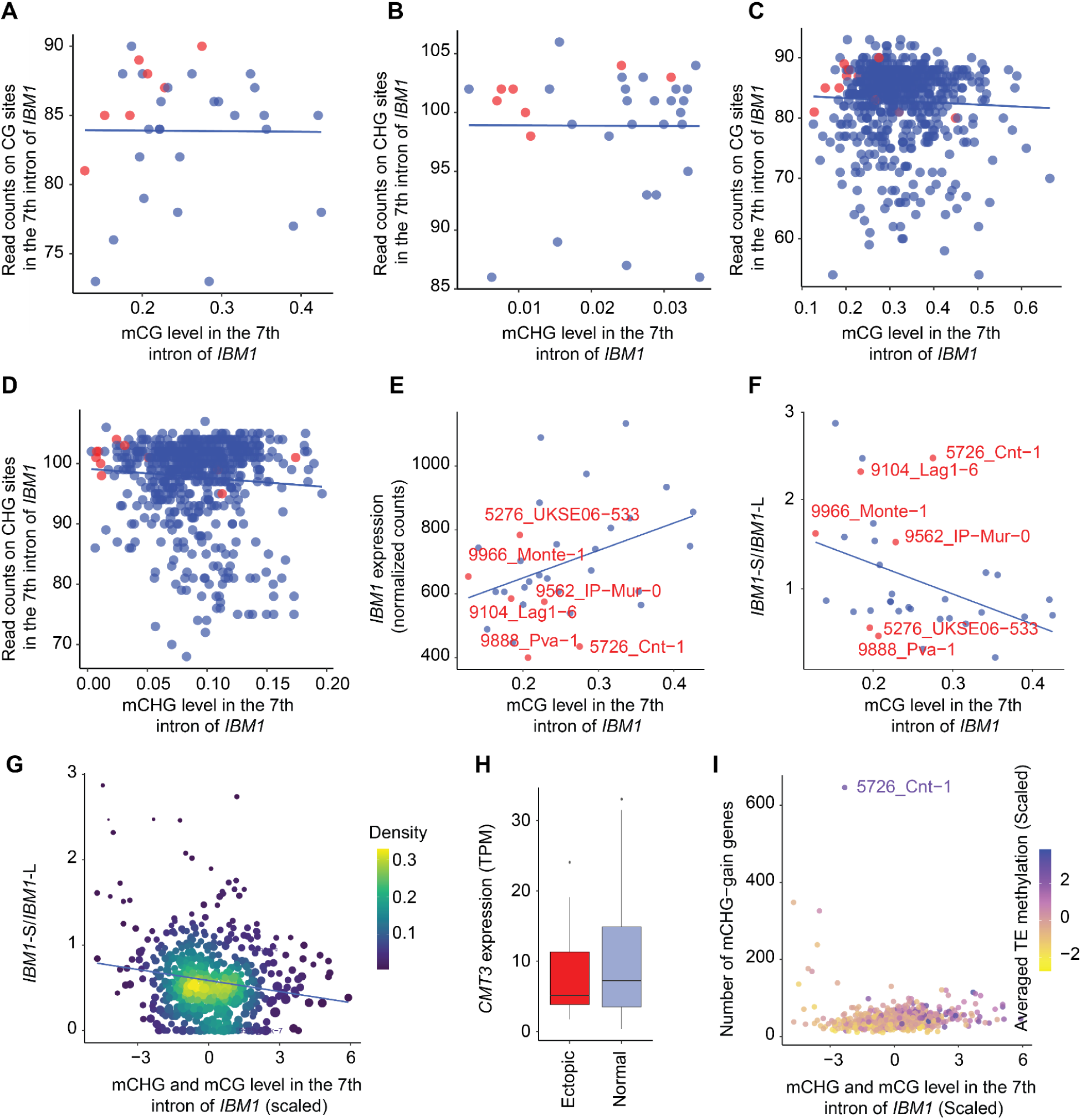
Features related to *IBM1* intronic methylation in *A. thaliana* natural accessions. **A.** Scatter plots (A-D) show the read counts on the *IBM1* intron region against the intron DNA methylation level of the long intron of *IBM1*. This plot shows the read counts on the CG sites of the *IBM1* intron region against the CG methylation level of the long intron of *IBM1*. Only accessions with mCHG level <0.035 in the long intron of *IBM1* were included. Accessions with a significant number (n≥120) of ectopic methylated genes are marked with red dots, whereas other accessions are represented with blue dots. **B.** Similar to A, except for the read count on the CHG sites against the CHG methylation level. **C.** Similar to A, except for including all natural accessions. **D.** Similar to B, except for including all natural accessions. **E.** The scatter plot shows the relationship between CG methylation levels in the long intron of *IBM1* and *IBM1* gene expression for accessions with low *IBM1* long intron mCHG level (<= 0.035). The top six accessions with the highest number of ectopic methylated genes are labeled by their name in red color. **F.** Similar to E, this scatter plot presents the CG methylation level in the long intron of *IBM1* plotted against the ratio of short to long isoform of *IBM1*. **G.** Similar to F, except for including all natural accessions. **H.** The box plot shows the distribution of *CMT3* gene expression by comparing accessions that rank in the top 12 for the highest number of ectopic methylated genes with all remaining accessions. **I.** The scatter plot shows the scaled CG and CHG methylation level in *IBM1’s* long intron against the number of genes with ectopic non-CG methylation in each natural accession. Accession Cnt-1 with the highest number (n=646) of ectopic methylated genes is labeled by its name. Each dot represents an accession and is colored by their averaged TE methylation level across the whole genome.

**Figure S2.**
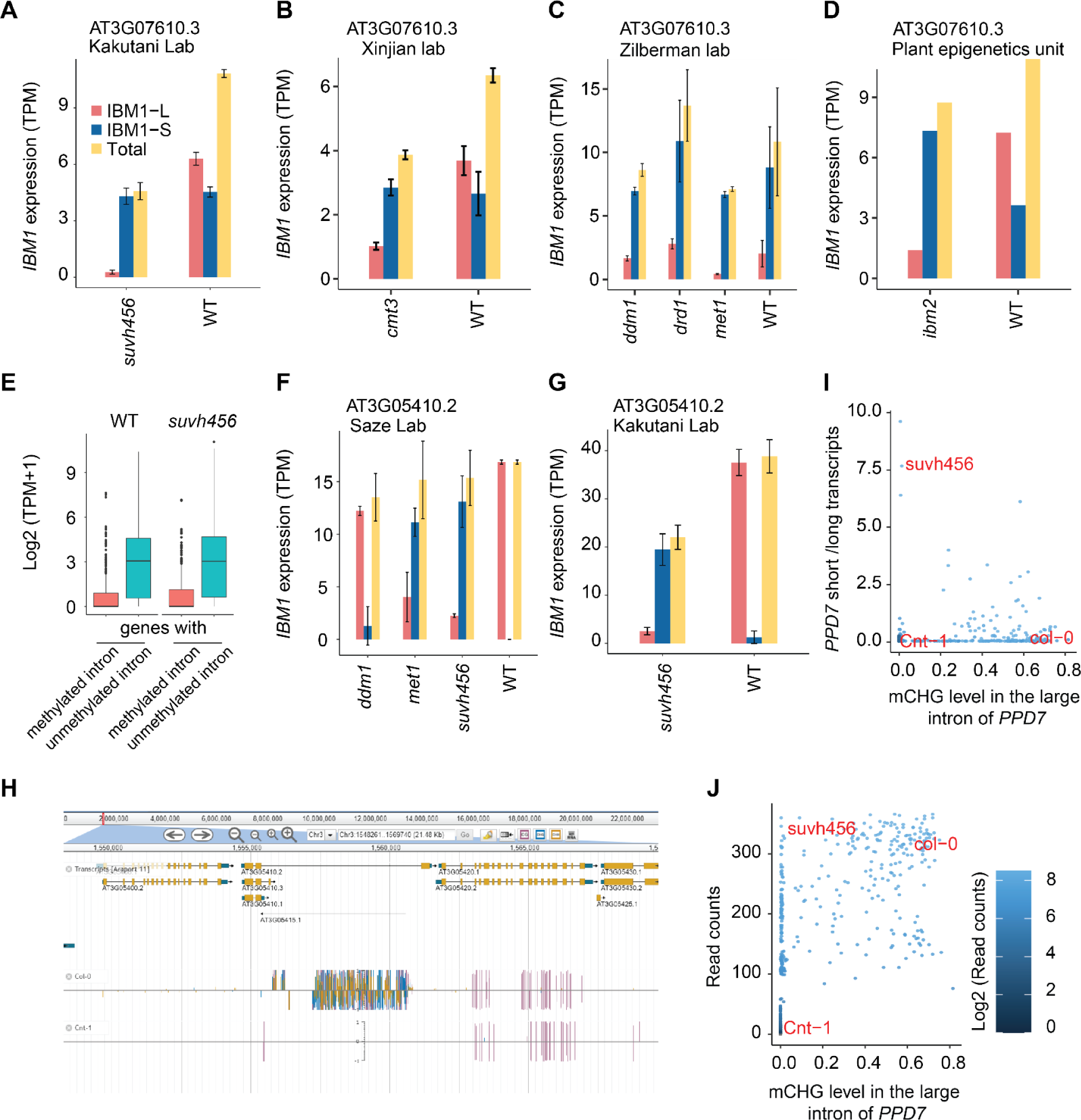
Features related to *IBM1* intronic methylation in *A. thaliana* mutant lines. **A.** The bar plot shows the expression levels of the long and short isoforms of *IBM1*, as well as their combined expression, in *suvh456* mutants, and Col-0 WT using RNA-seq data sourced from the Kakutani lab. **B.** Jianxi lab, **C.** Zilberman lab, **D.** Plant epigenetics unit. **E.** Genes featuring long introns (≥1kb) were categorized into two groups based on the presence or absence of CHG methylation. The box plot compares the expression distribution between these two groups. It displays the expression patterns of genes in both the wild type (right panel) and suvh456 mutant (left panel). **F.** the long and short isoforms of *PPD7* using RNA-seq data sourced from Saze lab, and **G.** Kakutani lab. **H.** A genome browser view shows the intron methylation pattern of *PPD7* in Col-0, which is lost in Cnt-1. **I.** The scatter plot shows the relationship between mCHG levels in the long intron of *PPD7* against the ratio of short to long isoform of *PPD7* for all natural accessions. The *suvh456* mutant, Cnt-1 and Col-0 are highlighted by their name in red color, while other accessions are represented by blue dots. **J.** The scatter plot shows the relationship between CHG methylation levels in the long intron of *PPD7* against the read counts on the CHG sites of long intron of *PPD7*. The color scheme is the same as H.

**Figure S3.**
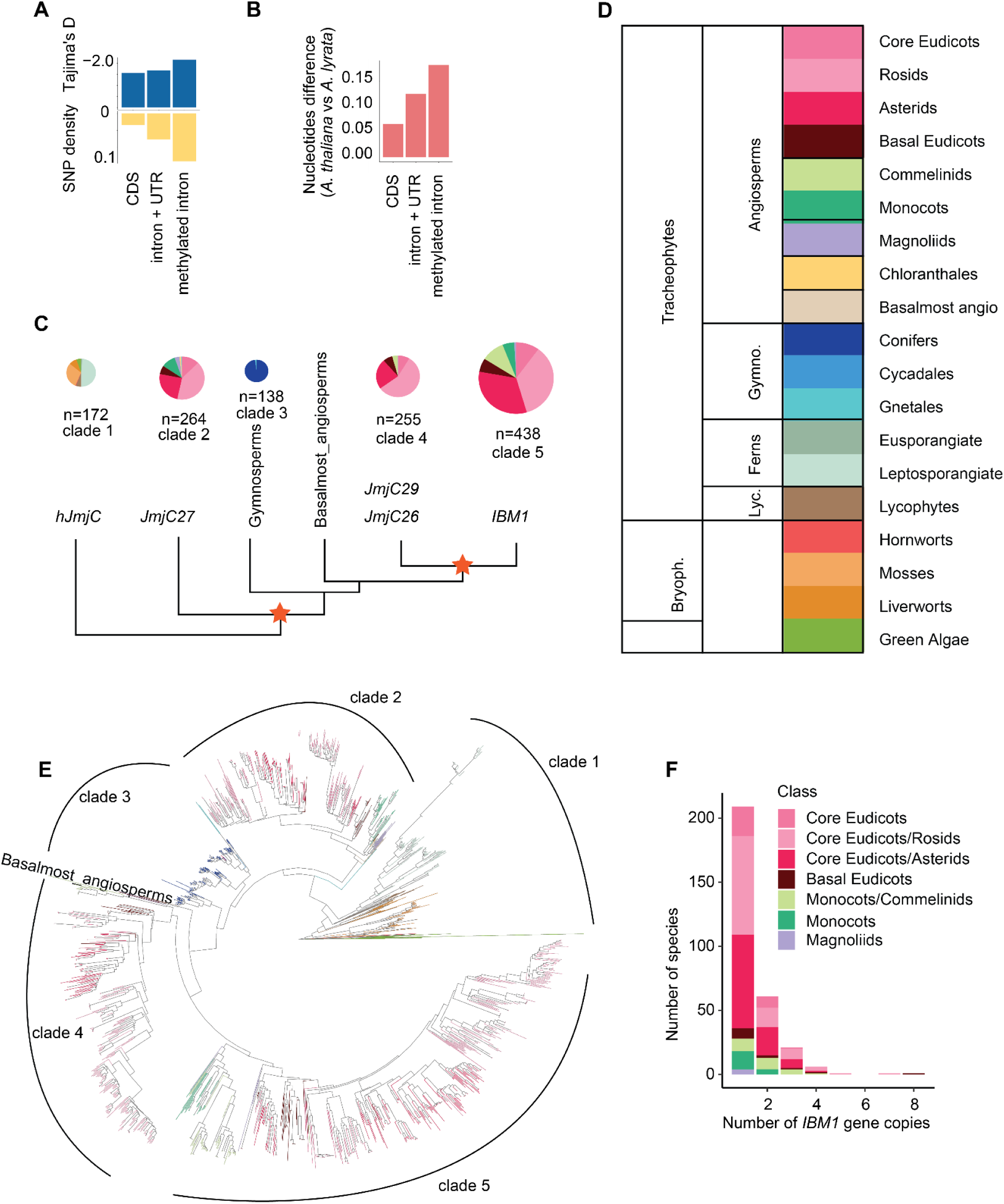
Phylogenetic relationships of JmjC family homologous genes across plants. **A.** The plot shows Tajima’s D and SNP density for *IBM1*, comparing its coding sequence (CDS), noncoding regions (inclusive of UTRs and introns not encompassed by non-CG methylation), and the region of methylated introns (denoted as m-intron). The SNP data for this analysis was sourced from the 1,001 Genomes Project. **B.** The plot illustrates the nucleotide differences, calculated as the proportion of mismatched bases, between *A. thaliana* and *A. lyrata*. These differences are shown separately for the DNA sequences in the coding sequence (CDS), non-coding regions, and the methylated intron region. **C.** The gene family tree of the JmjC domain is presented as a collapsed version, delineating six distinct subclades (also described in C). Pie charts, scaled according to the number of species, depict the species diversity within each clade. Two significant duplication events are highlighted: one shared by all angiosperms and gymnosperms, and another exclusive to all angiosperms except for the basalmost group (indicated by a star symbol). These events led to the diversification of various JmjC homologous genes, such as *JmjC27*, *JmjC26/29*, and *IBM1*. The tree is rooted in the clade that encompasses all green algae and liverwort species. **D.** The plot provides the color scheme for species classification as used in Fig. S3. **E.** The circular gene tree illustrates JmjC homologous genes organized into six distinct clades (the same as C), following the relationships of *A. thaliana* JmjC genes. These clades include: (1) homologous JmjC genes in green algae, bryophytes, and ferns; (2) *JmjC2*7; (3) JmjC genes in gymnosperms; (4) JmjC genes in basalmost angiosperms; (5) *JmjC26/29*; and (6) *IBM1*. Notably, the *JmjC26/29* and *IBM1* clades encompass all angiosperm species, with the exception of the basalmost angiosperms. This distribution is due to a duplication event that occurred just prior to the divergence of other angiosperms. **F.** The bar plot displays the distribution of the number of *IBM1* gene copies across all species within the *IBM1* gene clade.

**Figure S4.**
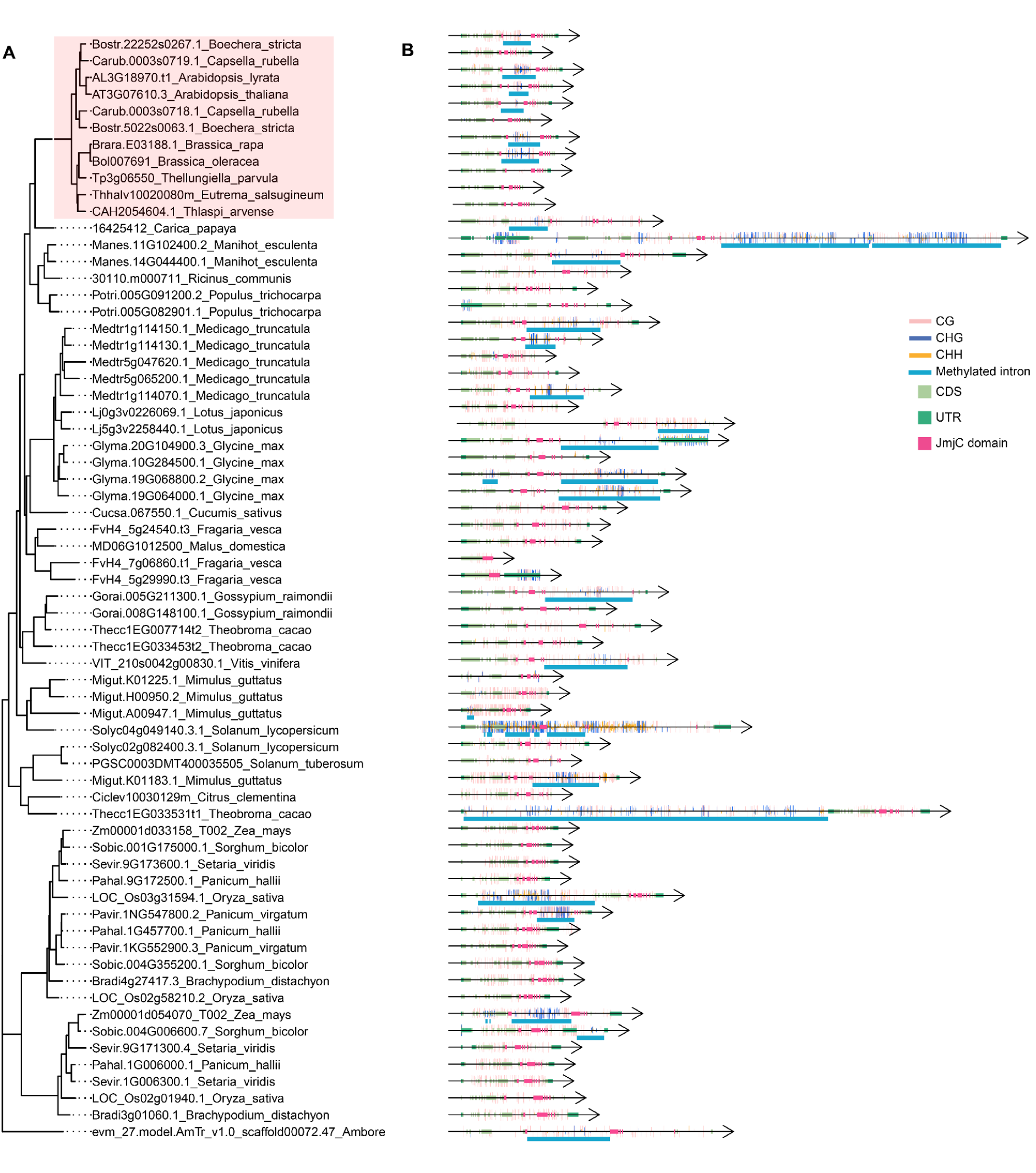
A complete version of the maximum likelihood gene tree that includes 65 *IBM1* orthologs. The gene tree has the same format and color scheme as Figure 3A.

**Figure S5.**
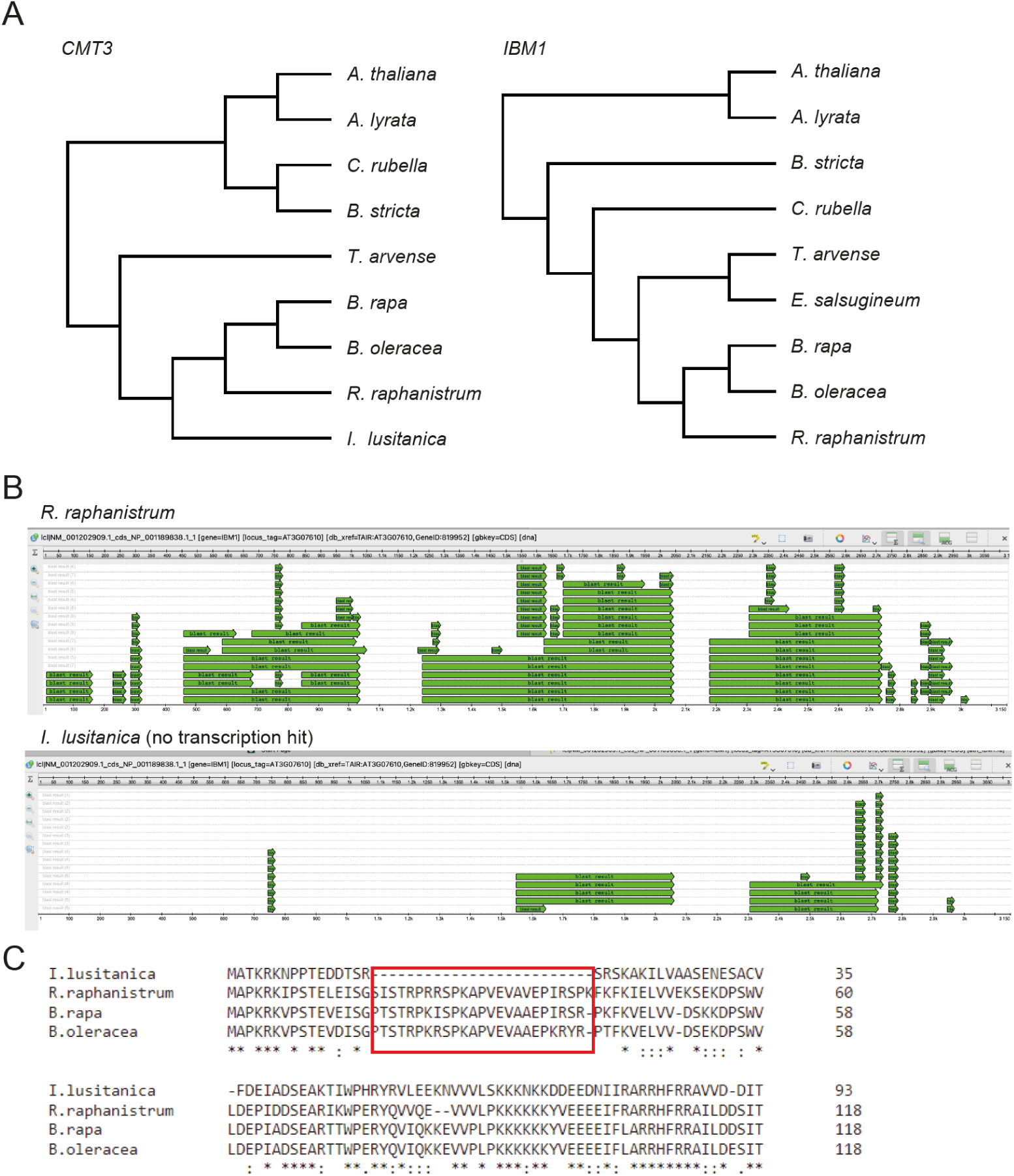
Identification of Brassicaceae species with potential loss or truncations of *IBM1* or *CMT3.* **A.** Ortholog analysis of *CMT3* and *IBM1* in the Brassicaceae species. **B.** De-novo assembly of *IBM1* with RNA-seq data. The assembled fragments of each species was blasted to A. thaliana IBM1 coding sequence. **C.** N-terminal deletion in CMT3 of *I. lusitanica*.

**Figure S6.**
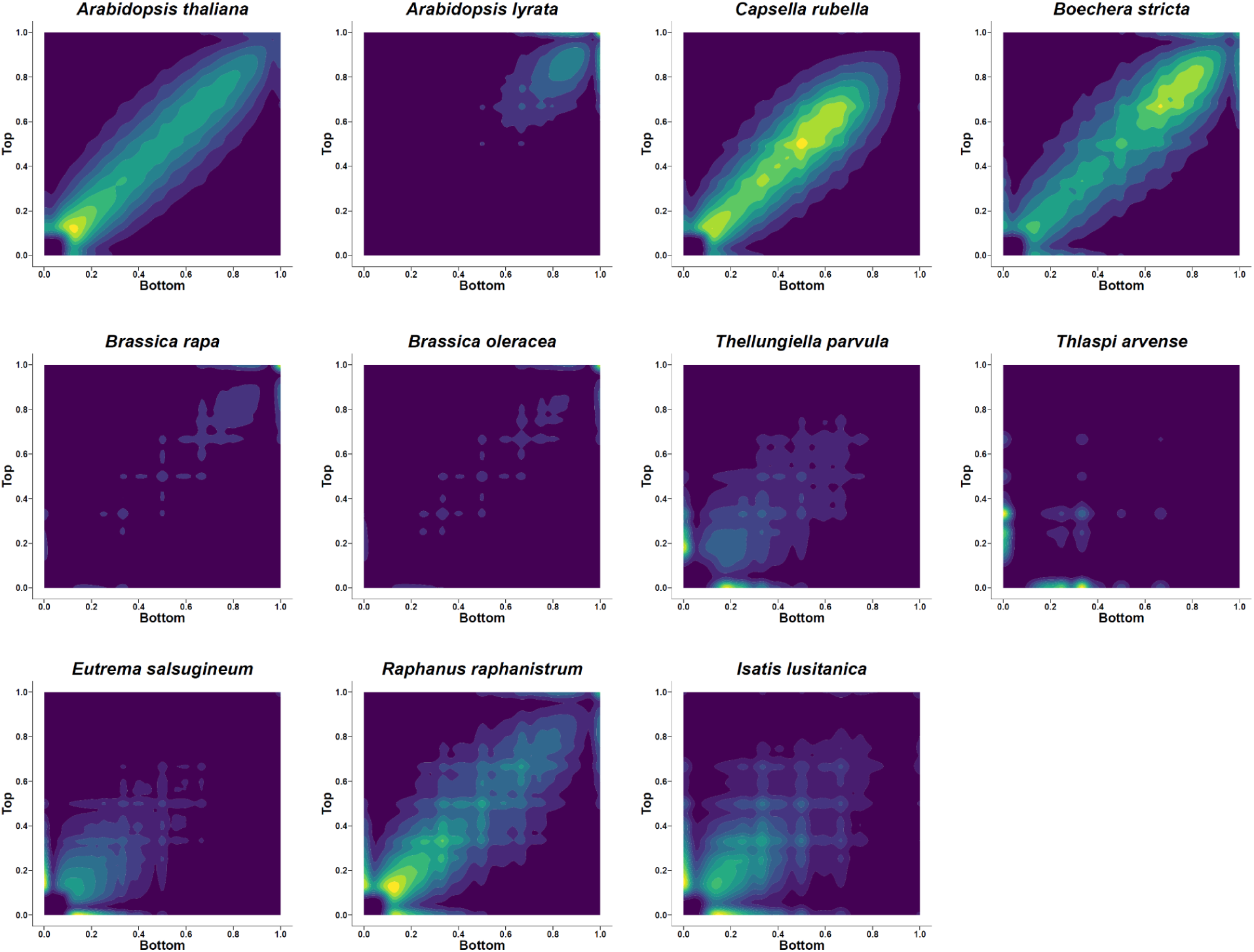
CWG methylation symmetry analysis in Brassicaceae species.

